# Widespread allelic heterogeneity in complex traits

**DOI:** 10.1101/076984

**Authors:** Farhad Hormozdiari, Anthony Zhu, Gleb Kichaev, Ayellet V. Segrè, Chelsea J.-T. Ju, Jong Wha J Joo, Hyejung Won, Sriram Sankararaman, Bogdan Pasaniuc, Sagiv Shifman, Eleazar Eskin

**Affiliations:** Department of Computer Science, University of California, Los Angeles, CA 90095, USA; Bioinformatics IDP, University of California, Los Angeles, CA 90095, USA; Cancer Program, The Broad Institute of Massachusetts Institute of Technology and Harvard University, Cambridge, MA 02142, USA; Neurogenetics Program, Department of Neurology, David Geffen School of Medicine, University of California Los Angeles, CA 90095, USA; Department of Pathology and Laboratory Medicine, University of California, Los Angeles, CA 90095, USA; Department of Human Genetics, University of California, Los Angeles, CA 90095, USA; Department of Genetics, The Institute of Life Sciences, The Hebrew University of Jerusalem, Jerusalem

## Abstract

Recent successes in genome-wide association studies (GWASs) make it possible to address important questions about the genetic architecture of complex traits, such as allele frequency and effect size. One lesser-known aspect of complex traits is the extent of allelic heterogeneity (AH) arising from multiple causal variants at a locus. We developed a computational method to infer the probability of AH and applied it to three GWAS and four expression quantitative trait loci (eQTL) datasets. We identified a total of 4152 loci with strong evidence of AH. The proportion of all loci with identified AH is 4-23% in eQTLs, 35% in GWAS of High-Density Lipoprotein (HDL), and 23% in schizophrenia. For eQTLs, we observed a strong correlation between sample size and the proportion of loci with AH (*R*^2^=0.85, P = 2.2e-16), indicating that statistical power prevents identification of AH in other loci. Understanding the extent of AH may guide the development of new methods for fine mapping and association mapping of complex traits.

## 1 Introduction

Genome-wide association studies (GWASs) have successfully identified many loci associated with various diseases and traits [1–4]. Unfortunately, interpreting the detected associated genes is challenging due to two facts: First, most of the associated variants fall in non-coding regions of the genome [5–10]. Second, for only a handful of GWAS loci a causal sequence variant was detected that underlies the trait or disease susceptibility. Therefore, it is challenging to identify the relevant genes, which is the first step to understanding the biological mechanisms of the disease.

Detecting the causal variants is complicated by the fact that most significant associated variants may not be causal, but instead be in linkage disequilibrium (LD) with unknown functional variants. In general, sequence variants, with respect to a trait, can be grouped into three categories. The first category is causal variants that have a biological effect on the trait and are responsible for the association signal. The second category is variants that are statistically associated with the trait due to high LD with the causal variants. The third category is variants that are not statistically associated with the trait and are not causal. Fine-mapping methods aim to distinguish between the two first categories (causal variants vs. correlated variants). One way to link the causal variant with a particular gene is by colocalization methods that determine whether a single variant is responsible for both variation in the trait and variation in expression of a gene at the same locus (expression quantitative trait loci, eQTLs).

Fine-mapping and colocalization methods are designed to identify the causal variant and the associated gene at a locus, but in many cases, they assume a single causal variant. In the presence of multiple causal variants, those fine-mapping and colocalization methods [11–13] will have a lower accuracy to detect the true causal variants and genes. Thus, a fundamental question is how many different causal variants are present in a locus

The presence of multiple causal variants, at the same locus that influence a particular disease or trait is known as allelic heterogeneity (AH). AH is very common for Mendelian traits. Clearly, many different mutations in the same gene may cause loss or gain of function leading to specific Mendelian disease. For example, approximately 100 independent mutations are known to exist at the cystic fibrosis locus [14], and even more are present at loci causing inherited haemoglobinopathies [15]. In contrast to Mendelian traits, the extent of AH at loci contributing to common, complex disease is almost unknown. Hermani et al. [16] have shown in their study the existence of multiple reproducible epistatic effects that influence gene expression. However, a recent study has shown that most of these epistatic effects can be explained by a third variant in that locus [17]. Thus, it is of utmost importance to detect loci that harbor AH in order to avoid considering them as epistatic interactions.

Identifying the number of causal variants in complex traits is difficult because of extensive LD and small effect sizes. The standard approach to identify AH is to use conditional analysis. In conditional analysis, the independent association of multiple SNPs is tested after conditioning on other SNPs, which are more significant. The conditional analysis lacks power because it requires multiple variants to have a strong effect and to be independently significant. When several associated variants are highly correlated, the conditional analysis will not detect multiple independent associations, but we cannot rule out the existence of AH. Thus, the extent of AH in complex traits is unknown. AH could substantially influence our ability to explain the missing heritability and to identify causal genes.

We developed and applied a new method to quantify the number of independent causal variants at a locus responsible for the observed association signals in GWAS. Our method is incorporated into the CAusal Variants Identification in Associated Regions (CAVIAR) software [18]. The method is based on the principle of jointly analyzing association signals (i.e., summary level Z-scores) and LD structure in order to estimate the number of causal variants. Our method computes the probability of having multiple independent causal variants by summing the probability of all possible sets of SNPs for being causal. We compared results from our method to results produced using the standard conditional method (CM) [19], which tests for independent association of a variant after conditioning on its significantly associated neighbors. Using simulated datasets, we illustrate that CAVIAR tends to outperform CM. We observed a very low false positive rate for CAVIAR to detect loci with AH even when the true causal variant is not included in our dataset. We applied CAVIAR to both eQTL and GWAS datasets. Our results indicate that in the Genotype-Tissue Expression (GTEx) dataset [20] 4-23% of eGenes harbor AH. We observed a high correlation between the portions of loci with AH and sample size. In addition, we replicated a significant fraction of the loci with AH in three other existing eQTL studies. We also applied CAVIAR to three GWAS datasets, schizophrenia (SCZ) [3], high-density lipoprotein (HDL) [21], and major depression disorder (MDD) [22], where we observed 23%, 35%, and 50% of loci (respectively) with strong evidence for AH.

## 2 Results

### 2.1 Overview of identifying allelic heterogeneity

Our method utilizes the shape of marginal statistics (statistics obtained from GWAS such as z-score) and the patterns of LD to detect whether or not a locus harbors AH. In Figures 1A and 1B, we illustrate a simple example with no AH. However, we can guess that the region shown in Figure 1C harbors AH; here, we observe two high independent peaks in the region. Unfortunately, detecting AH is less intuitive in regions that are more complicated.

**Figure 1.**
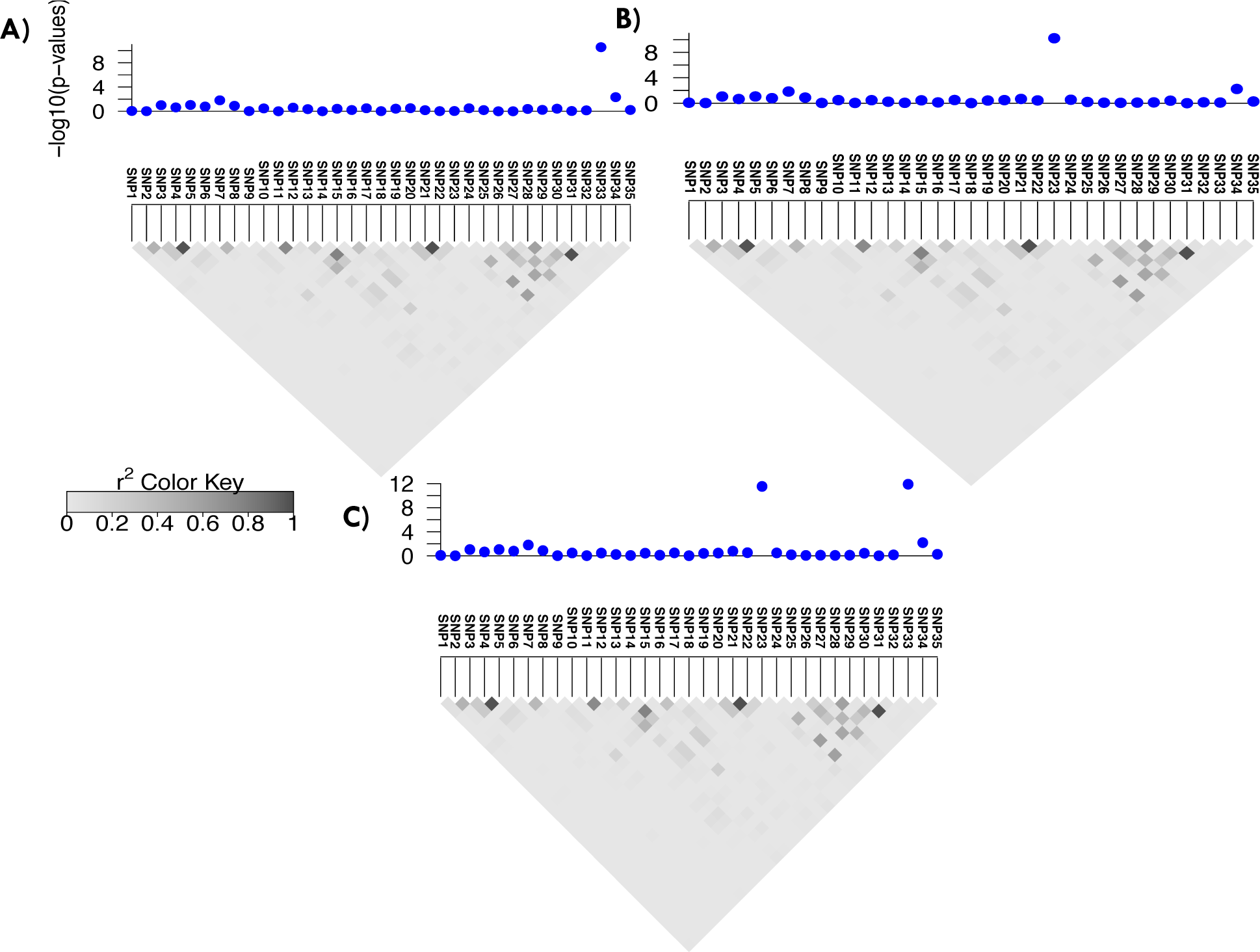
Overview of CAVIAR for Detecting Allelic Heterogeneity Regions. Panels (A) and (B) indicate the marginal statistics for a locus where we have implanted one causal variant. In panel (A), SNP23 is causal and in panel (B), SNP33 is causal. Panel (C) is the same locus where both SNP23 and SNP33 are causal. In these figures, the x-axis is the negative logarithm of the p-values for each locus to indicate the strength of the marginal statistics. The grey triangle below each figure indicates the LD pattern. Each square indicates the correlation between two variants and the magnitude of the correlation is shown by the color intensity of the square. The darker the square the higher the correlation between two variants.

The input to our method is the LD structure of the locus and the marginal statistics for each variant in the locus. The LD between each pair of variants is computed from genotyped data or is approximated from HapMap or 1000G data [23, 24]. We use the fact that the joint distribution of the marginal statistics follows a multivariate normal distribution (MVN) [18, 25–27] to compute the posterior probability of each subset of variants being causal, as described below. Then, we compute the probability of having i independent causal variants in a locus by summing the probability of all possible subsets of size i (sets that have i causal variants). We consider a locus to be AH when the probability of having more than one independent causal variant is more than 80%.

We would like to emphasize that only using summary statistics and LD information is insufficient for differentiating tightly linked variants. For example, if two variants are in perfect pairwise LD (correlation of 1), it is impossible to detect with just the summary statistics whether only one of the variants is causal or both are causal. Therefore, our estimates are just a lower bound on the amount of loci with AH for a given complex trait.

### 2.2 CAVIAR is more accurate than existing methods

In order to assess the performance of our method, we generated simulated data sets. We used HAPGEN2 [28], a widely used software, to generate a case-control study using the European population obtained from the 1000G. Then, we implanted one or two causal variants in a region and generated simulated phenotypes using the linear mixed model (described in the Method section).

After generating the phenotypes, we performed a t-test to generate the marginal statistics for each variant. We applied CAVIAR to all the simulated data sets to detect loci that harbor AH. We generated two datasets. In the first datasets, we set the non-centrality parameter (NCP) to have 10%, 30%, 50% or 70% power to detect the causal variants. In the second datasets, we set the NCP of the causal variants to have 20%, 40%, 60%, or 80% power to detect the causal variants. We compared our results with the conditional method (CM). We use false positive (FP) and true positive (TP) as metrics for comparison. FP indicates the fraction of loci with one causal variant that are incorrectly detected as loci with AH. TP indicates the fraction of loci with AH that are correctly detected. We found that our method has higher TP compared to CM for the same FP rate. Figure 2 and Supplementary Figure 1 illustrate the ROC curves for the first and second simulated datasets.

**Figure 2.**
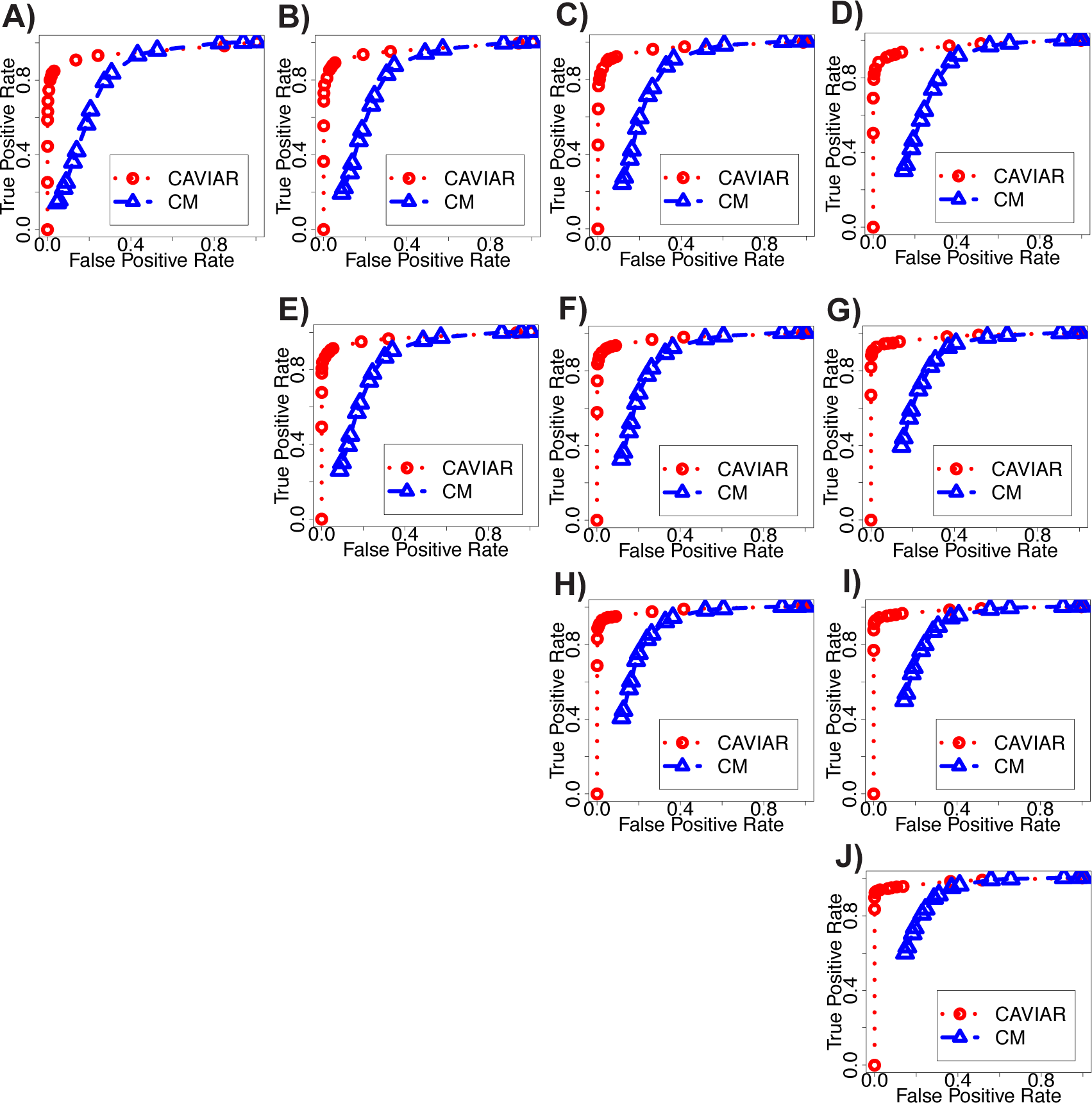
ROC curve for our method (CAVIAR) and CM. We implant one causal variant to compute the false positive (FP) rate. FP indicates loci that harbor one causal variant; however, these loci are detected as AH. We implant two causal variants to compete the true positive (TP) rate. TP indicates loci that harbor AH and are detected correctly. We range the NCP such that the power at the causal variant is 20%, 40%, 60%, and 80% at the genome significant level 10^−8^. In (A), (B), (C), and (D) NCP for the first causal SNP is set to 20% power. In (A), (B), (C), and (D) NCP for the second causal SNP is set to 20%, 40%, 60%, and 80%, respectively. In (E), (F), and (G) NCP for the first causal SNP is set to 40% power. In (E), (F), and (G) NCP for the second causal SNP is set to 40%, 60%, and 80%, respectively. In (H) and (I) NCP for the first causal SNP is set to 60% power. In (H) the NCP of the second causal SNP is set to 60% power and in (I) the NCP of the second causal SNP is set to 80%. in (J) the NCP for both causal SNPs are set to 80% power.

**Figure 3.**
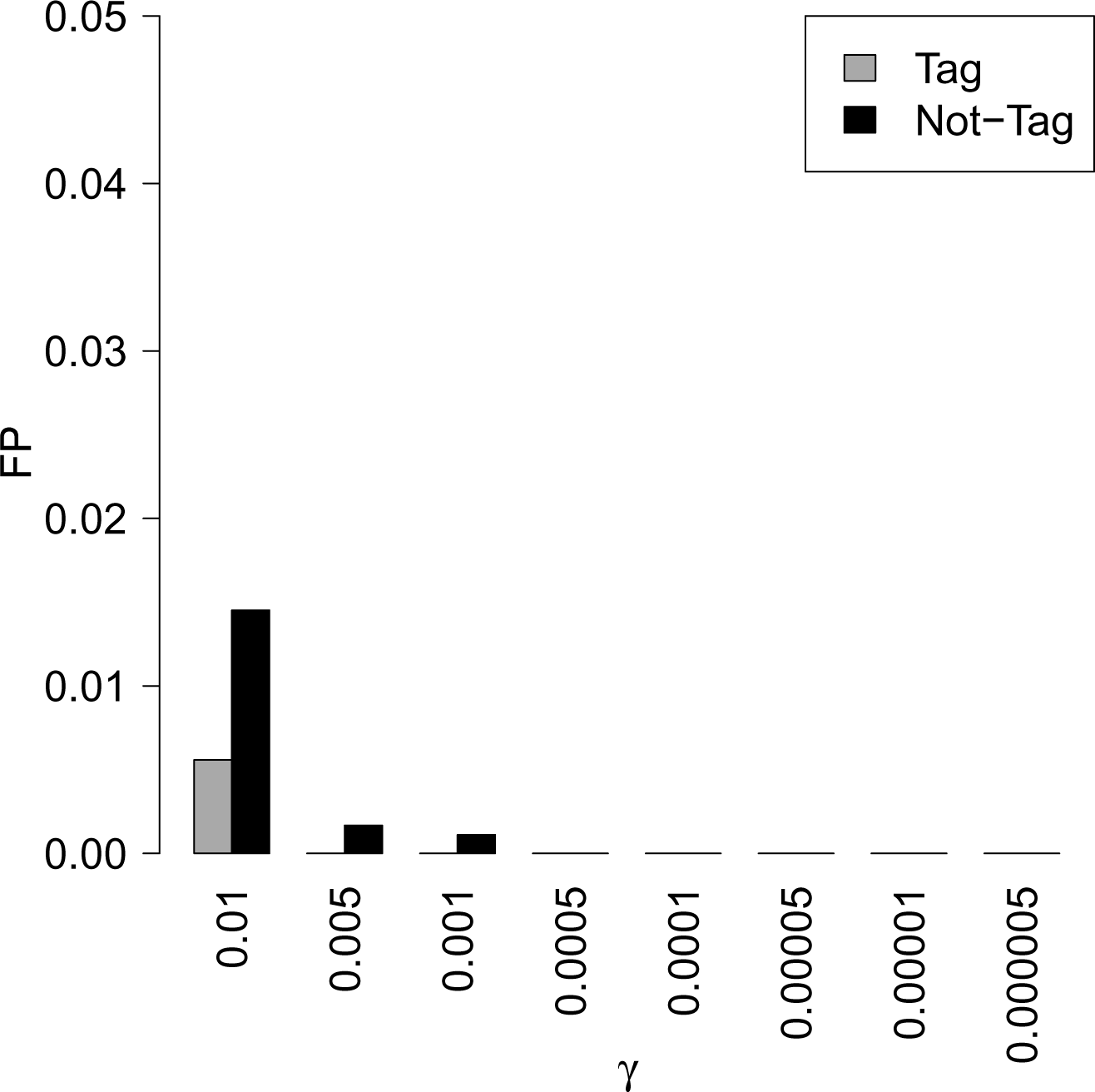
CAVIAR has low FP even when the true causal variant is not collected (untagged). Thus, most loci that are detected by CAVIAR to harbor AH are most probably true. X-axis indicates the prior probability of causal variant (γ). We set γ to 0.01, 0.005, 0.001, 0.0005, 0.0001, 0.00005, 0.000001, and 0.000005.

### 2.3 CAVIAR has a low false positive even when the causal variant is not included

In the previous section, we show CAVIAR has an extremely low FP and a high TP rate in detecting loci with AH. In the simulated data, the causal variant was included. However, in real datasets we cannot guarantee that information will exist for all the causal variants. One possible reason for detecting a locus as AH could be that the actual causal variant is not included or tagged in the data. In this section, we show that loci detected by CAVIAR with AH are rarely due to the fact that the actual causal variant is not included.

We simulate datasets where we implant one causal variant in a locus and generate marginal statistics in a method similar to the previous section. Next, we remove the causal variants from our analysis and use the remaining variants in the locus as an input to CAVIAR. We observe the FP is extremely low (FP < 0.015), even when the causal variant was not included in the analyzed data (see Figure 2). Our conclusion is that CAVIAR may fail to detect AH in some loci, but a very small proportion of loci where we detected AH do not harbor AH.

### 2.4 CAVIAR is robust to different input parameters

There are two main input parameters to CAVIAR, excluding the summary statistics of a locus: the prior probability that a variant is causal (γ) and LD structure. The LD structure can be computed from raw genotypes when it is available. However, in most datasets, we do not have access to the raw genotypes. Thus, we approximate the LD utilizing the 1000G [23, 24] or HapMap [29] dataset.

We simulated marginal statistics in a way similar to previous sections. Then, we vary γ among 0.01, 0.005, 0.001, 0.0005, 0.0001, 0.00005, 0.000001, and 0.000005. We observed that for different values of γ, CAVIAR tends to have extremely low FP (FP ≤ 0.001) and high TP. The result for this experiment is shown in Supplementary Figure 2.

In the next experiment, we want to investigate the effect of using misspecified LD structures. We simulated the marginal statistics by utilizing the LD structure (LD matrix) obtained from HAPGEN2. After, we simulate the marginal statistics, we generate a misspecified LD by adding standard Gaussian noise, *N*(0,τ), to each element of the original LD matrix. We use the simulated marginal statistics and the misspecified LD as an input to CAVIAR. We simulated 10,000 loci that harbor one causal variant to compute the FP, and we simulated 10,000 loci that harbor two causal variants to measure TP. We vary the variance of the Gaussian noise (τ) between 0.01, 0.02, 0.05, 0.1, 0.2, 0.25, and 0.3. In this experiment, we observe low FP and high TP for CAVIAR results. The result for this experiment is shown in Supplementary Figure 3.

### 2.5 CAVIAR accurately detects the number of causal variants in a locus when all the variants are collected

In the previous sections, we have shown that CAVIAR is accurate in detecting loci that harbor AH. An additional benefit of our new method is that it can accurately detect the number of causal variants in a locus. We simulated the phenotypes similar to the previous sections. In these experiments, we implanted one, two, and three causal variants in a locus. We set the NCP of the causal variant such that the statistical power is 20%, 30%, 40%, 50%, 60%, 70%, and 80%.

As CAVIAR provides probability values for different number of causal variants, we consider a locus to have *i* independent causal variants where the probability of having *i* causal variants is the maximum probability for different numbers of causal variants. In the case of CM, the number of causal variants in a locus is equivalent to the number of conditional steps that we perform until the p-value of all variants is higher than 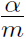 (Bonferroni correction) where *m* is the total number of variants in a locus and α is 0.05. We compute recall rate as the fraction of simulations where any methods correctly predicated the number of causal variants in a locus. In Figure 4, we plot the recall rate of CAVIAR and CM in detecting the number of causal variants. We observe that CAVIAR has much higher recall rate in detecting the true number of causal variants in a locus than CM.

**Figure 4.**
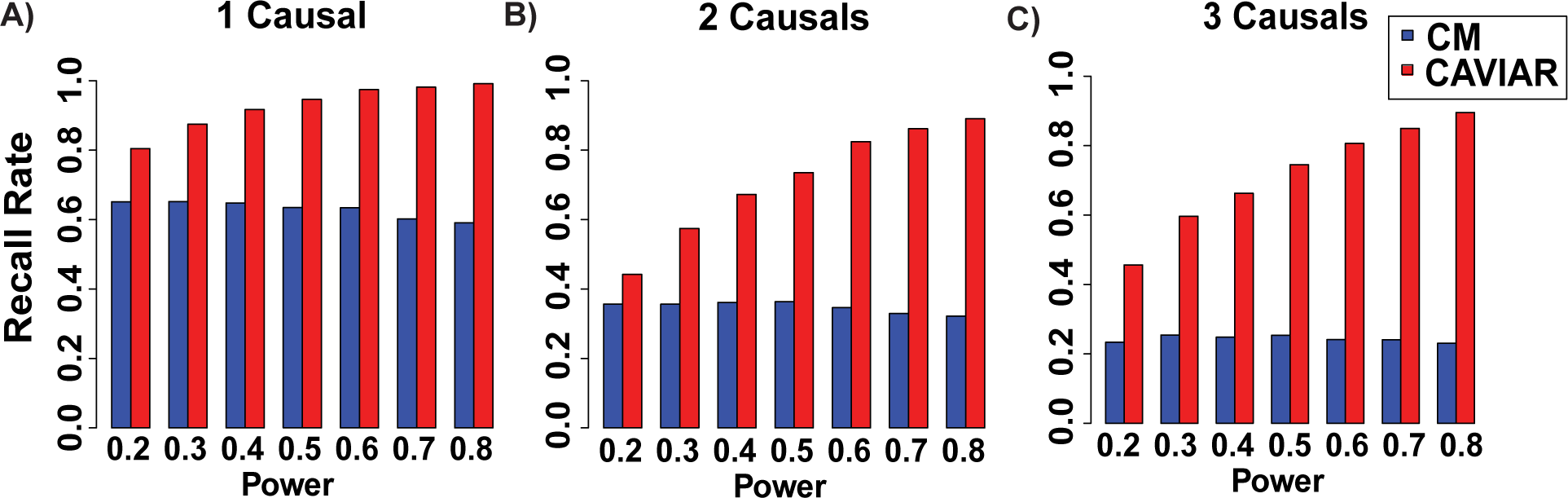
CAVIAR is more accurate than CM to detect the number of causal variants. The x-axis is the power of causal variants and the y-axis is the accuracy to detect the number of causal variants in a locus. We implanted one, two, and three causal variants. (A-C) Recall rate of each method for different number of causal variants, (A) one causal variant (B) two causal variants (C) three causal variants. We vary the statistical power to detect the causal variant among 0.2, 0.3, 0.4, 0.5, 0.6, 0.7, and 0.8.

### 2.6 CAVIAR distinguishes between epistatic interaction and allelic heterogeneity

It is possible to incorrectly detect a locus with AH due to the epistatic interaction in that locus. In this section, we utilize simulated data to illustrate that CAVIAR rarely detects AH in loci where the true genetics architecture is epistasis. We simulated different datasets where we implanted epistatic interactions between two randomly selected variants in a locus. Then, we generated simulated phenotypes using the linear additive model (described in the Method section). We vary the number of individuals in each dataset among 500, 1000, 1500, 2000, 2500, and 3000. In addition, for each dataset, we vary the effect size among 0.01, 0.02, 0.03, 0.04, 0.05, 0.06, 0.07, 0.08, 0.09, 0.1, and 0.2. For each simulated dataset, we simulated 5,000 different marginal statistics. CAVIAR has extremely low false positive in these experiments (see Figure 5). In Figure 5, we show the results for the effect size of 0.04, 0.05, 0.06, 0.07, 0.08, 0.09, 0.1, and 0.2. The results for effect size of 0.01, 0.02, and 0.02 are not shown as the FP is zero for these values. As a result, CAVIAR only detects a small fraction of epistatic interactions as AH. It is worth mentioning that the amount of epistatic interactions for different traits is very low. Thus, these results indicate the loci detected by CAVIAR that harbor AH are not artifacts of epistatic interaction.

**Figure 5.**
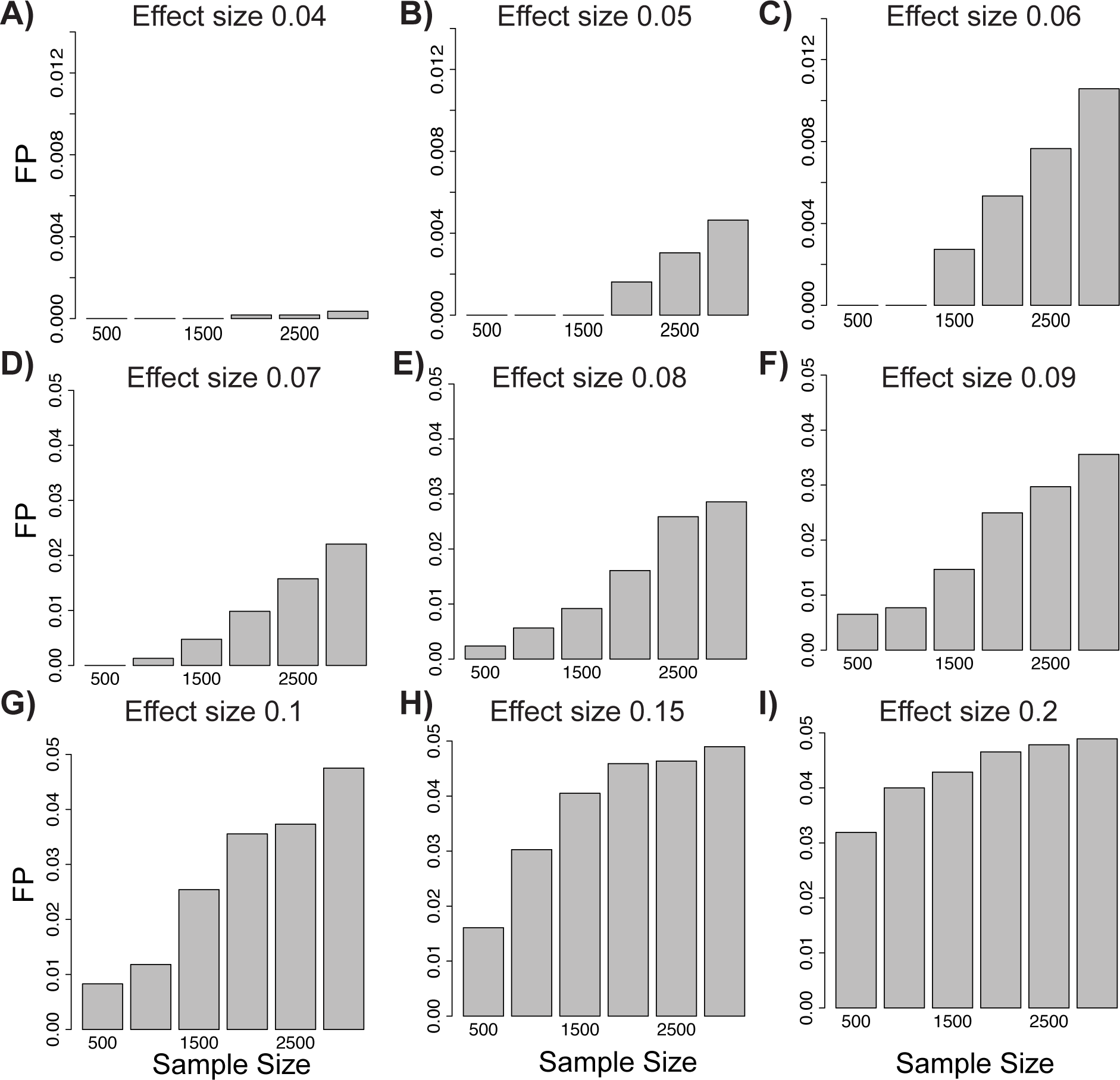
CAVIAR distinguishes between epistatic interaction and allelic heterogeneity. The x-axis is the sample size that we vary between 500, 1000, 1500, 2000, 2500, and 3000 individuals and the y-axis is the false positive (FP) rate. We simulated datasets where we have epistatic interaction and compute the FP as the number of cases where CAVIAR incorrectly detects these loci to harbor AH. (A-I) illustrate the FP for different effect sizes of the epistatic interaction.

### 2.7 Prevalence of allelic heterogeneity in eQTL datasets

We used four datasets to examine the extent of AH in eQTL datasets. We utilize version v6p of data, which consists of 44 tissues. For each tissue, we have around 22,000 genes (probes). For each gene in each tissue, we have access to the marginal statistics for the cis-eQTL analysis, which we obtained from the GTEx project [20] and genotype data that is used to compute the LD pattern for each gene. Then, we filtered out genes lacking at least one significant SNP. We set the significant cut-off threshold to a p-value of 10-5. Genes that have a significant cis-eQTL SNP are known as eGene. We applied our method to detect AH loci only to eGene. We found that 4%-23% of the eGenes show evidence for AH (with probability > 80%) (Figure 6, Table 1). In addition, we applied the CM to the same set of eGenes. We observed that 50%-80% of loci detected by the CM to have AH were also detected by CAVIAR (see Table 1).

**Figure 6.**
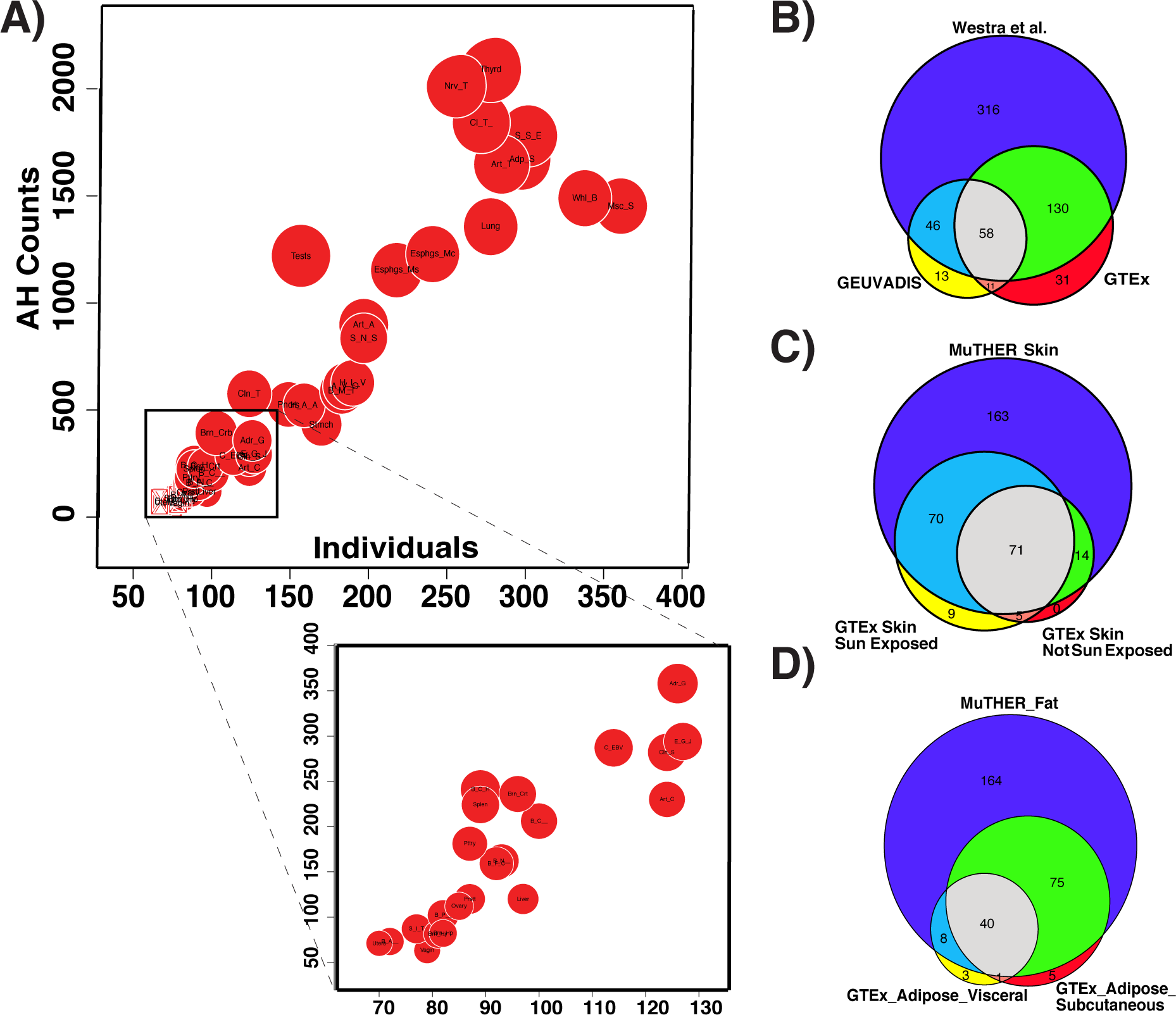
Levels of allelic heterogeneity in eQTL studies. (A) Linear relationship between the amount of AH and sample size. Each red circle indicates a different type of tissue from the GTEx dataset. The size of each red circle is proportional to the number of genes that harbor a significant eQTL (eGenes). (B-D) Significant overlap between AH estimations for different eQTL datasets, shown for (B) blood (P=7.9e-97), (C) skin (P=4.9e-63), and (D) adipose (P=1.1e-69).

**Table 1.**
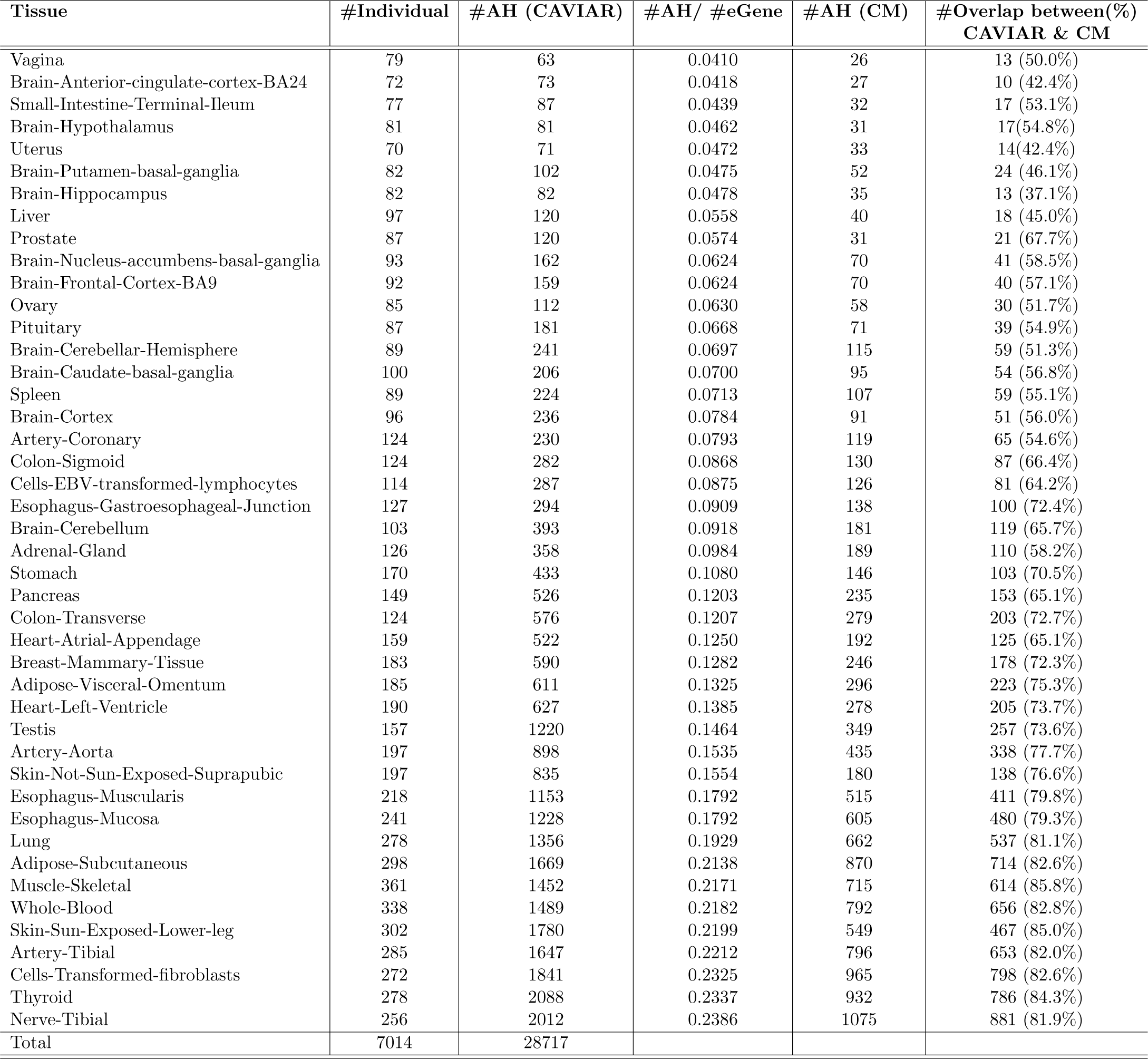
List of 44 tissues in GTEx. Tissues are sorted based on the number of samples. #Individual indicates the number of samples for each tissue. #AH (CAVIAR) is the number of loci detected by CAVIAR that harbor AH. #AH(CAVIAR)/#eGene is the fraction of eGenes that are detected to harbor AH. #AH(CM) is the number of loci detected by conditional method (CM) that harbor AH.

| Tissue | #Individual | #AH (CAVIAR) | #AH/ #eGene | #AH (CM) | #Overlap between(%)<br>CAVIAR & CM |
| --- | --- | --- | --- | --- | --- |
| Vagina | 79 | 63 | 0.0410 | 26 | 13 (50.0%) |
| Brain-Anterior-cingulate-cortex-BA24 | 72 | 73 | 0.0418 | 27 | 10 (42.4%) |
| Small-Intestine-Terminal-Ileum | 77 | 87 | 0.0439 | 32 | 17 (53.1%) |
| Brain-Hypothalamus | 81 | 81 | 0.0462 | 31 | 17(54.8%) |
| Uterus | 70 | 71 | 0.0472 | 33 | 14(42.4%) |
| Brain-Putamen-basal-ganglia | 82 | 102 | 0.0475 | 52 | 24 (46.1%) |
| Brain-Hippocampus | 82 | 82 | 0.0478 | 35 | 13 (37.1%) |
| Liver | 97 | 120 | 0.0558 | 40 | 18 (45.0%) |
| Prostate | 87 | 120 | 0.0574 | 31 | 21 (67.7%) |
| Brain-Nucleus-accumbens-basal-ganglia | 93 | 162 | 0.0624 | 70 | 41 (58.5%) |
| Brain-Frontal-Cortex-BA9 | 92 | 159 | 0.0624 | 70 | 40 (57.1%) |
| Ovary | 85 | 112 | 0.0630 | 58 | 30 (51.7%) |
| Pituitary | 87 | 181 | 0.0668 | 71 | 39 (54.9%) |
| Brain-Cerebellar-Hemisphere | 89 | 241 | 0.0697 | 115 | 59 (51.3%) |
| Brain-Caudate-basal-ganglia | 100 | 206 | 0.0700 | 95 | 54 (56.8%) |
| Spleen | 89 | 224 | 0.0713 | 107 | 59 (55.1%) |
| Brain-Cortex | 96 | 236 | 0.0784 | 91 | 51 (56.0%) |
| Artery-Coronary | 124 | 230 | 0.0793 | 119 | 65 (54.6%) |
| Colon-Sigmoid | 124 | 282 | 0.0868 | 130 | 87 (66.4%) |
| Cells-EBV-transformed-lymphocytes | 114 | 287 | 0.0875 | 126 | 81 (64.2%) |
| Esophagus-Gastroesophageal-Junction | 127 | 294 | 0.0909 | 138 | 100 (72.4%) |
| Brain-Cerebellum | 103 | 393 | 0.0918 | 181 | 119 (65.7%) |
| Adrenal-Gland | 126 | 358 | 0.0984 | 189 | 110 (58.2%) |
| Stomach | 170 | 433 | 0.1080 | 146 | 103 (70.5%) |
| Pancreas | 149 | 526 | 0.1203 | 235 | 153 (65.1%) |
| Colon-Transverse | 124 | 576 | 0.1207 | 279 | 203 (72.7%) |
| Heart-Atrial-Appendage | 159 | 522 | 0.1250 | 192 | 125 (65.1%) |
| Breast-Mammary-Tissue | 183 | 590 | 0.1282 | 246 | 178 (72.3%) |
| Adipose-Visceral-Omentum | 185 | 611 | 0.1325 | 296 | 223 (75.3%) |
| Heart-Left-Ventricle | 190 | 627 | 0.1385 | 278 | 205 (73.7%) |
| Testis | 157 | 1220 | 0.1464 | 349 | 257 (73.6%) |
| Artery-Aorta | 197 | 898 | 0.1535 | 435 | 338 (77.7%) |
| Skin-Not-Sun-Exposed-Suprapubic | 197 | 835 | 0.1554 | 180 | 138 (76.6%) |
| Esophagus-Muscularis | 218 | 1153 | 0.1792 | 515 | 411 (79.8%) |
| Esophagus-Mucosa | 241 | 1228 | 0.1792 | 605 | 480 (79.3%) |
| Lung | 278 | 1356 | 0.1929 | 662 | 537 (81.1%) |
| Adipose-Subcutaneous | 298 | 1669 | 0.2138 | 870 | 714 (82.6%) |
| Muscle-Skeletal | 361 | 1452 | 0.2171 | 715 | 614 (85.8%) |
| Whole-Blood | 338 | 1489 | 0.2182 | 792 | 656 (82.8%) |
| Skin-Sun-Exposed-Lower-leg | 302 | 1780 | 0.2199 | 549 | 467 (85.0%) |
| Artery-Tibial | 285 | 1647 | 0.2212 | 796 | 653 (82.0%) |
| Cells-Transformed-fibroblasts | 272 | 1841 | 0.2325 | 965 | 798 (82.6%) |
| Thyroid | 278 | 2088 | 0.2337 | 932 | 786 (84.3%) |
| Nerve-Tibial | 256 | 2012 | 0.2386 | 1075 | 881 (81.9%) |
| Total | 7014 | 28717 |  |  |  |

The number of eGenes detected in a tissue depends on the statistical power to detect a significant variant associated with the gene expression. The statistical power is highly correlated to the number of samples for that tissue. We hypothesized that there might also exist correlation between the sample size and the number of loci with AH. Indeed, we observed that the proportion of eGenes with AH for each tissue is in a linear relationship with the sample size (*R*^2^=0.85, P = 2.2e-16). This result indicates that statistical power prevents the identification of AH at other loci.

To check the reproducibility of the AH detection, we compared the results from GTEx blood data with results from two other blood eQTL studies: GEUVADIS [30] and Wester et al. (2013) [31]. We tested the overlap between genes with AH for skin and adipose tissues based on the GTEx [20] and MuTHER [32] datasets. We only considered eGenes that are common between the studies. In all comparisons, we observed a high reproducibility rate for the detection of AH in blood (Figure 6B, P=7.9e-97), skin (Figure 6C, P=4.9e-63), and adipose (Figure 6D, P=1.1e-69) tissues.

### 2.8 Prevalence of allelic heterogeneity in GWAS datasets

To measure the level of AH in a human quantitative trait, we applied our method to a GWAS of High-Density Lipoprotein (HDL)[21]. Out of 37 loci, 13 (35%) showed evidence for AH with probability ≥ 80% (see Supplementary Table 1). We also studied the results of GWASs focused on two psychiatric diseases: major depression disorder (MDD) [22] and schizophrenia (SCZ) [3]. For MDD, we found evidence for AH at one of two loci. For SCZ, we identified 25 loci out of 108 (23%) with high probability of AH (see Supplementary Table 2). One example of AH in SCZ is the locus on chromosome 18 that includes the *TCF4* gene (Figure 7A). The locus contains multiple associated SNPs that are distributed in different LD blocks (Figure 7B). According to our analysis, there are three or more causal variants in this locus with high probability (Figure 7C) (for similar results in other loci, see Supplementary Figures 4-39 for HDL and Supplementary Figures 40-167 for SCZ).

**Figure 7.**
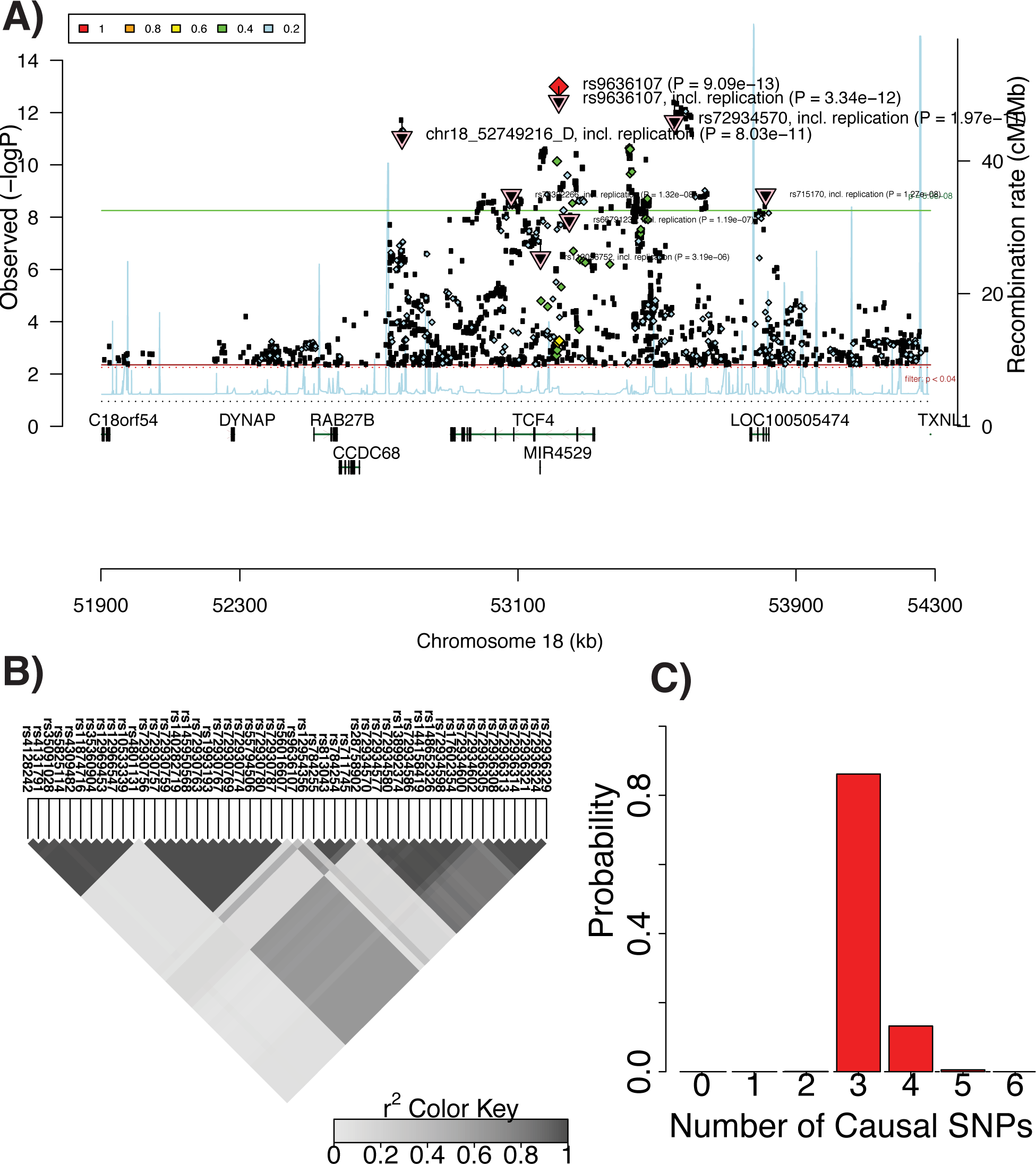
Allelic heterogeneity in the TCF4 locus associated with schizophrenia. (A) Manhattan plot obtained from Ricopili (http://data.broadinstitute.org/mpg/ricopili/) consists of all the variants in a 1Mbp window centered on the most significant SNP in the locus (rs9636107). This plot indicates multiple significant variants that are not in tight LD with the peak variant. (B) LD plot of the 50 most significant SNPs showing several distinct LD blocks. (C) Histogram of the estimated number of causal variants.

## 3 Methods

### 3.1 Joint Distribution of Observed Statistics in Standard GWAS

In this section, we provide a brief description of statistical tests that are performed in GWAS and the joint distribution of computed marginal statistics. These statistics are used as an input to CAVIAR. We consider, we perform GWAS on a quantitative traits for n individuals. Let *Y* be a vector of (*n* × 1) where *y_i_* indicates phenotypic value for the *i*-th individual. Moreover, we genotype all the individuals for all the *m* variants. Let 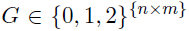 be a matrix of genotypes for all the individuals where *g_ij_* is the minor allele count for the *i*-th individual at the *j*-th variant. We standardize the minor allele count for each variant to have mean of zero and variance one. We use *X_j_* to indicate the standardized minor allele count for all the individuals at the *j*-th variant. We have 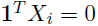 and 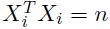 as the genotypes are standardized.

We assume that phenotypic values follow the linear model and the *c* variant is the causal variant. Thus, we have:

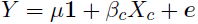

where *μ* is the phenotypic mean value, *β_c_* is the effect size of the variant *X_c_*, **1** is a (*n* × 1) vector of ones, and ***e*** is a (*n* × 1) vector to model the environment contribution and error in the measurement. In this model, we assume the error are i.i.d. and follows a normal distribution with mean of zero and variance of 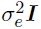, where ***I*** is a (*n* × *n*) matrix of identity and *σ_e_* is the variance scalar of ***e***. From the model explained above, we have 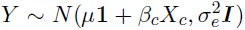. We use the maximum likelihood to compute the estimated *β_c_* and *σ_e_* which are denoted by 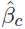 and 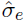. Thus, we have:

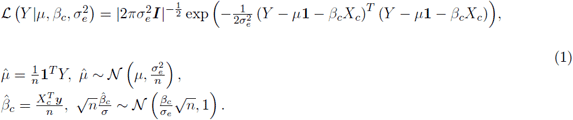

The association statistic for SNP *c*, denoted by *S_c_*, follows a non-central t-distribution, which is the ratio of a normally distributed random variable to the square root of an independent chi-squared distributed random random variable. As shown in previous works, if the number of individuals are large enough [18], we can assume the marginal statistics follows a normal distribution with mean equal to 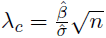 and variance of one,

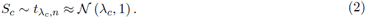

We assume that variant *i* is correlated with the causal variant *c* and the correlation between the two variants are *r* where 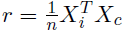. Then, the marginal statistics estimated at variant *i* is equal to the marginal statistics for the causal variant that is scaled by *r*. We compute the covariance of the marginal statistics between two variants where the LD between the two variants are *r*.

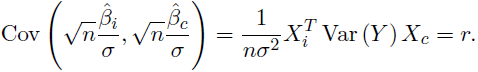

We compute the joint distribution of the marginal statistics for two variants *i* and *j* as follow:

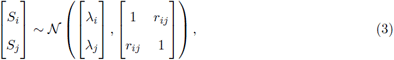
where 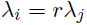 if the variant *i* is causal and the variant *j* is not causal or 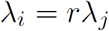 if the variant *j* is causal and the variant *i* is not causal.

### 3.2 Computing the Likelihood of Causal Configuration

We can extend the joint distribution of marginal statistics for two variants to more general case. Assume we have *m* variants and the pair-wise correlation between each variant is denoted by Σ. Let 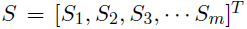 be the vector of marginal statistics obtained for each variant. The joint distribution of the marginal statistics for all the m variants is computed as follow:

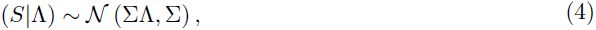

where Λ is (*m* × 1) is a vector of normalized effect sizes and λ_i_ is the normalized effect size of the *i*-th variant. We introduce a new parameter *C* that indicates the causal status of each variant. Each variant can have two possible causal status. We have *c_i_* = 1 if the *i*-th variant is causal and *c_i_* = 0 if the *i*-th variant is not a causal variant. We define a prior probability on the vector of effect size Λ for a given causal status using a multivariate normal distribution,

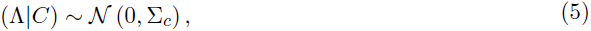

where 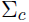 is a (*m* × *m*) matrix. The off diagonal elements of 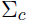 are set to zero. The diagonal elements are set to σ or zero. We set the *i*-th diagonal element to σ if the *i*-th variant is causal and we set *i*-th diagonal element to zero if the *i*-th variant is not causal. Thus, the joint distribution follows a multivariate normal distribution,

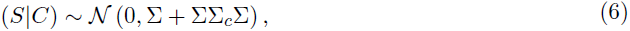

### 3.3 Computing the Number of Independent Causal Variants in a Locus

In this section, we provide the formula to compute the probability of having *i* causal variants in a locus. We compute the probability of having *i* causal variants in a locus by summing over all the possible causal configurations where only *i* variants are causal. Let *N_c_* indicates the number causal variants in a locus. We have,

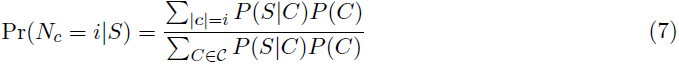

where *P*(*C*) is the prior on the causal configuration *C, **C*** is the set of all possible causal configurations including the configuration all the variants are not causal and |*C*| indicates the number of causal variants in the causal configuration *C*. The numerator in the above equation considers all possible causal configurations that have *i* causal variants, and the denominator is a normalization factor to ensure the probability definition holds.

In this paper, we use a simple prior for a causal status. We assume the probability of a variant to be causal is independent from other variants and the probability of a variant to be causal is γ. Thus, we compute the prior probability as, 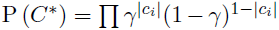. We utilize different values for γ as shown in Supplementary Figure 2. In our experiment, we set γ to 0.001 [33–35]. It is worth mentioning, although we use a simple prior for our model, we can incorporate external information such as functional data or knowledge from previous studies. As a result, we can have variant-specific prior where *γ_i_* indicates the prior probability for the *i*-th variant to be causal. Thus, we can extend the prior probability to a more general case, 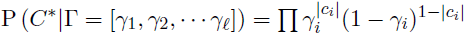.

### 3.4 Reducing the Computational Complexity for Computing the Likelihood of a Causal Status

Unfortunately, the time complexity to compute the likelihood of a causal status using Equation (6) is *O*(*m*^3^). In this section, we provide a speed up process that reduces the time complexity to *O*(*m*^2^*k*) where *k* is the number of causal variants for a causal status. The number of causal variants is smaller than the total number of variants in a locus (*k* << *m*). Thus, we manage to speed up the likelihood computation by a factor of 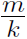.

According to Equation (6) to compute the likelihood of a causal status, we require to compute the following quantity:

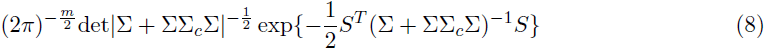

where det|.| denotes the determinant of a matrix. First, we speed up the exponential part. Thus, we have:

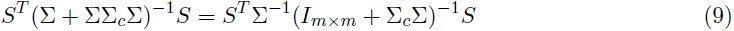

where 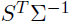 is independent from the causal status and can be computed once and used many times. As a result, the expensive computational part is to compute 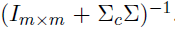. We set elements of *U* and *V* such that 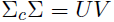. Let *α_i_* indicate the index of i-th causal variant. We set elements of *V* as follows: 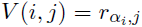. Let 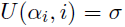 and we set the rest of elements of *U* to zero. Thus, we have:

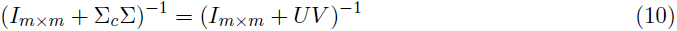

We use Woodbury matrix identity formula to compute the inverse. We have:

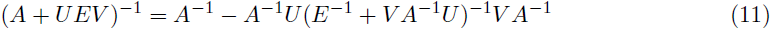

We set *A* to 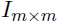 and *E* to 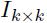, then the left side of Woodbury matrix identity formula converts to 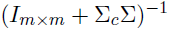. From the right side of Woodbury matrix identity formula, we have:

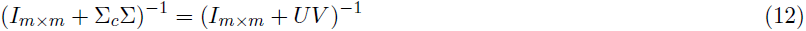

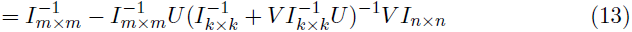

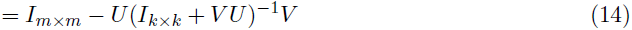

where 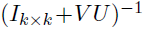 requires inverting a (*k* × *k*) matrix that is much faster than inverting a (*m* × *m*) matrix.

In similar way, the naive method to compute det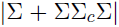 requires *O*(*n*^3^). We utilize the Sylvester’s determinant identity that is as follows:

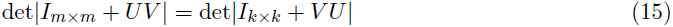

Thus, instead of computing the determinant of a (*m* × *m*) matrix, we can compute the same value by computing the determinant of a (*k* × *k*) matrix. We have:

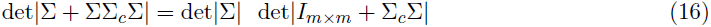

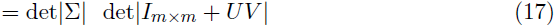

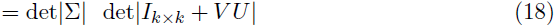

In above equation, we can compute det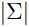 once and use it for different causal statuses. In addition, the computational complexity of det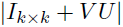 is 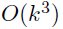. Thus, the time complexity to compute det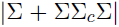 is 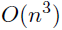.

### 3.5 Conditional Method (CM)

A standard method to detect allelic heterogeneity (AH) is the conditional method (CM). In CM, we identify the SNP with most significant association statistics. Then, conditioning on that SNP, we re-compute the marginal statistics of all the remaining variants in the locus. We consider a locus to have AH when the re-computed marginal statistics for at least one of the variants is more significant than a predefined threshold. Similarly, we consider a locus to not have AH when the re-computed marginal statistics of all variants fall below the predefined threshold. The predefined threshold is referred to as the stopping threshold for CM. This standard method can be applied to either summary statistics or individual level data. GCTA-COJO [19] performs conditional analysis while utilizing the summary statistics.

When applying CM to individual level data, we re-compute the marginal statistics by performing linear regression where we add the set of variants that are selected as covariates. We utilize the LD between the variants, which we obtain from a reference dataset, when applying CM to summary statistics data. In this case, we re-compute the marginal statistics for the ith variant as follows:

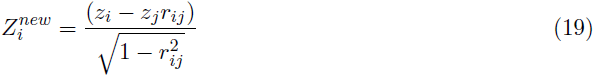

when we have selected the jth variant as causal. Let *z_i_* indicate the marginal statistics for the ith variant and *r_ij_* the genotype correlations between the ith and jth variants.

### 3.6 Datasets

**<u>Genotype-Tissue Expression (GTEx):</u>** We obtained the summary statistics for GTEx eQTL dataset (Release v6p, dbGaP Accession phs000424.v6.p1). We estimated the LD structure using the available genotypes in the GTEx dataset. We considered 44 tissues and applied our method to all eGenes, genes that have at least one significant eQTL, in order to detect loci that harbor allelic heterogeneity.

**<u>Genetic European Variation in Disease (GEUVADIS):</u>** We obtained the summary statistics of blood eQTL for 373 European individuals from the GEUVADIS website. We approximated LD structure from the 1000G CEU population. We applied our method to the 2954 eGenes in GEUVADIS to detect AH loci.

**<u>Multiple Tissue Human Expression Resource (MuTHER):</u>** We obtained the summary statistics from the MuTHER website. We utilized the skin and fat (adipose) tissues. We then approximated LD from the 1000G CEU population. We obtained 1433 eGenes for skin and 2769 eGenes for adipose.

**<u>High-Density Lipoprotein Cholesterol (HDL-C):</u>** We used the High-Density Lipoprotein Cholesterol (HDL-C) trait [21]. We only considered the GWAS hits, which are reported in a previous study. We applied ImpG-Summary [36] to impute the summary statistics with 1000G as the reference panel. We identified 37 loci that have at least one causal variant. Following common protocol in fine-mapping methods, we assumed at least one causal variant. Then, we applied our method to each locus.

**<u>Psychiatric diseases:</u>** We analyzed the recent GWAS on major depression disorder and schizophrenia. The major depression disorder study has 2 and the schizophrenia study has 108 loci identified to contain at least one significant variant. We utilized the summary statistics provided by each study and approximated the LD using the 1000G CEU population.

### 3.7 Data simulation

**<u>Simulated data with no epistasis interaction:</u>** We first simulated genotypes using HAPGEN2[28], where we utilized the 1000G CEU population as initial reference panels. Then, we simulated phenotypes using the Fisher's polygenic model, where the effects of causal variants are obtained from the normal distribution with a mean of zero. Let Y indicate the phenotypes and X indicate the standardized genotypes. In addition, *β* is the vector of effect sizes where *β_i_* is the effect of the i-th variant. Thus, we have:

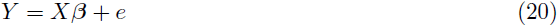
where e models the environment and measurement noise. Under the Fisher's polygenic model, the effect size of the causal variants is obtained from 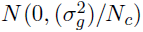, where *N_c_* is the number of causal variants and *σ_g_* is the genetic variation. In addition, the effect size for variants that are non-causal is zero. We set the effect size in order to obtain the desired statistical power. We implanted one, two, or three causal variants in our simulated datasets.

We use false positive (FP) and true positive (TP) as metrics to compare different methods. FP indicates the fraction of loci that harbor one causal variant and are incorrectly detected as loci that harbor AH. TP indicates the fraction of loci that harbor AH and are correctly detected.

**<u>Simulated data with epistasis interaction:</u>** We simulated the genotypes similar to the case where we have no epistasis interaction, which is mentioned above. Then, we simulated phenotypes using the following model:

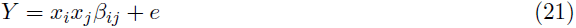
where *x_i_* and *x_j_* indicate the standardized genotypes for ith and jth variants, respectively. Moreover, *β_ij_* is the epistasis interaction effect size. We set *β_ij_* such that we obtained the desired heritability for the simulated phenotype. Then, we computed the marginal statistics for each variant utilizing the linear regression single variant testing.

## 4 Discussion

We have proposed a novel probabilistic method to detect loci with AH. Our results show that our method is more accurate than existing methods. One of the main benefits of our method is that it only requires summary statistics. Summary statistics of a GWAS study or eQTL study are widely available; thus, our method is applicable to most existing datasets. We have shown that AH is widespread and more common than previously estimated in complex traits, both in GWASs and eQTL studies. Since our method is influenced by statistical power and uncertainty induced by LD, the proportions of loci with AH detected in this study are just a lower bound on the true amount of AH. Thus, our study suggests that many, and maybe even most, loci are affected by AH.

Our results highlight the importance of accounting for the presence of multiple causal variants when characterizing the mechanism of genetic association in complex traits. Falling to account for AH can reduce the power to detect true causal variants, and can explain the limited success of fine mapping of GWASs. Similarly, attempts to explain GWAS using eQTLs data should be more successful with methods that assume that some loci may include multiple causative variants (e.g. eCAVIAR [37] and RTC [38]).

One of the limitations of our method is that we assume that the observed marginal statistics are corrected for the population using PCA-based methods. Recently, linear mixed models (LMM) [39–44] have become a popular correction for population structures that have cryptic relationships. Thus, the current version of our method is not applicable to summary statistics that have been corrected for population structure using LMM. However, we have shown in our previous work that the same statistical model can be extended to incorporate the summary statistics that have been corrected for population structure using LMM. Unfortunately, in this case, the study's raw genotypes and phenotypes should be available in order to perform the desired analysis.

In summary, we have developed a method to detect the presence of AH in loci of complex traits. We show that while the method may fail to detect AH in some loci, the false positive rate is very low. Thus, when our method detects a locus to have AH with a high probability, the prediction is very reliable. Since the amount of AH detected in our study is just a lower bound on the number of loci with AH, we suggest that AH is widespread in complex traits.

## 5 Acknowledgments

FH, JWJJ, and EE are supported by National Science Foundation grants 0513612, 0731455, 0729049, 0916676, 1065276, 1302448, 1320589 and 1331176, and National Institutes of Health grants K25-HL080079, U01-DA024417, P01-HL30568, P01-HL28481, R01-GM083198, R01-ES021801, R01-MH101782 and R01-ES022282. EE is supported in part by the NIH BD2K award, U54EB020403. AVS is supported by a contract (HHSN268201000029C) to the Laboratory, Data Analysis, and Coordinating Center (LDACC) at The Broad Institute, Inc. SS was supported in part by NIH grant R00-GM 111744-03. GK is supported by the Biomedical Big Data Training Program (NIH-NCI T32CA201160). We acknowledge the support of the NINDS Informatics Center for Neurogenetics and Neurogenomics (P30 NS062691).

## 6 Web Resources

CAVIAR is available http://genetics.cs.ucla.edu/caviar/

GTEx dataset (Release v6, dbGaP Accession phs000424.v6.p1) is available at http://www.gtexportal.org.

GEUVADIS dataset is available at ftp://ftp.ebi.ac.uk/pub/databases/microarray/data/experiment/GEUV/E-GEUV-1/analysis_results/.

MuTHER dataset is available at http://www.muther.ac.uk/Data.html.

Blood eQTL dataset is available at http://genenetwork.nl/bloodeqtlbrowser/.

